# Genetic engineering of carbon monoxide dehydrogenases produces distinct autotrophic phenotypes in *Clostridium autoethanogenum*

**DOI:** 10.64898/2026.02.27.708534

**Authors:** Kurshedaktar Majibullah Shaikh, Kristina Reinmets, Pratik Rajendra Pawar, Clara Vida G. C. Carneiro, Kaspar Valgepea

## Abstract

Acetogens are promising microbes for sustainable biomanufacturing but improving acetogen gas fermentation requires efficient conversion of CO and CO_2_ into fuels and chemicals. Carbon monoxide dehydrogenase (CODH) enzymes couple carbon fixation to energy conservation in acetogens and serve as potential regulatory modules for tuning autotrophic metabolism. Intriguingly, the model-acetogen *Clostridium autoethanogenum* lost its unique truncation in the bifunctional CODH (*acsA*), essential for autotrophy, during autotrophic adaptive laboratory evolution while obtaining superior phenotypes. Additionally, protein expression of the monofunctional CODH *cooS1* is high and conditionally-regulated in *C. autoethanogenum*. Here, we genetically engineered CODHs in *C. autoethanogenum* by replacing the stop codon in *acsA* with leucine (strain Leu_SNP) or serine (Ser_SNP), and deleting *cooS1* (Δ*cooS1*). Phenotyping in autotrophic batch and chemostat cultures revealed altered growth profiles and significant redistribution of carbon and redox flows in SNP strains, whereas Δ*cooS1* showed moderate and condition-dependent effects. Surprisingly, structural modelling identified no conformational differences between wild-type and mutant AcsA proteins. While transcriptomics showed limited transcriptional changes in Δ*cooS1*, it suggested potential transcriptional adjustments linked to reduced robustness and altered product profile of Leu_SNP. Our results demonstrate the impact of CODHs on autotrophy and offer targets for rational engineering of acetogen cell factories.

## INTRODUCTION

Rising energy demands and waste accumulation from a growing population have escalated environmental problems, largely due to fossil fuel combustion and insufficient waste recycling. Advances in renewable energy technologies are facilitating the decarbonisation of energy production. Meanwhile, developments in microbial engineering are further enabling biotechnological processes to replace fossil-based chemicals and use waste feedstocks for fuel and chemical production. Gas fermentation, utilising C1-fixing bacteria, offers a promising sustainable solution for recycling C1 gases (CO, CO◻, CH◻ ) from industrial waste gases and gasified waste into value-added products.^1,2^ *Clostridium autoethanogenum*, a model gas-fermenting acetogen, has been successfully utilised for the production of ethanol industrially, and acetone and isopropanol at pilot scale.^3,4^ *C. autoethanogenum* uses the most energy-efficient CO_2_-fixation pathway – the Wood-Ljungdahl pathway (WLP) – for fixing carbon oxides into acetyl-CoA^5^ and can tolerate high levels of CO (>50%)^6^ that are lethal to most microbes and inhibitory to many acetogens.^7–9^

The WLP enzymes in *C. autoethanogenum* are expressed from a C1-fixing gene cluster comprising of 16 genes associated with autotrophic growth.^10^ Carbon monoxide dehydrogenase (CODH) is one of the key enzymes responsible for *C. autoethanogenum* robust growth on CO, which catalyses the reversible oxidation of CO to CO_2_ using ferredoxin (Fd) as the electron carrier. ^10 11^, *C. autoethanogenum* encodes three CODH isoforms: CAETHG_1620-1621 (*acsA;* LABRINI_08025), CAETHG_3005 (*cooS1;* LABRINI_15015), and CAETHG_3899 (*cooS2;* LABRINI_19440), with

*acsA* being essential for autotrophic growth on CO and CO_2_+H_2_.^12^ Notably, *acsA* in the wild-type *C. autoethanogenum* JA1-1 strain features a TGA stop codon at position 401 (full-length 631 amino acids), splitting the gene into two fragments (CAETHG_1620 and CAETHG_1621), while other relevant acetogens primarily have serine at this position (Table 1). Truncation in this key C1-fixation gene is especially intriguing, although the full-length translational product has been observed at low levels^12^ with cysteine at the stop codon position.^11^ Interestingly, during autotrophic adaptive laboratory evolution (ALE) of *C. autoethanogenum* JA1-1, the TGA stop codon was mutated to leucine (TTA) in the isolate termed LAbrini, which showed superior growth characteristics and contained four more mutations.^13^ Therefore, phenotyping strains with mutations replacing this stop codon would improve our understanding of AcsA, thus also CODHs and acetogen metabolism.

**Table 1.**
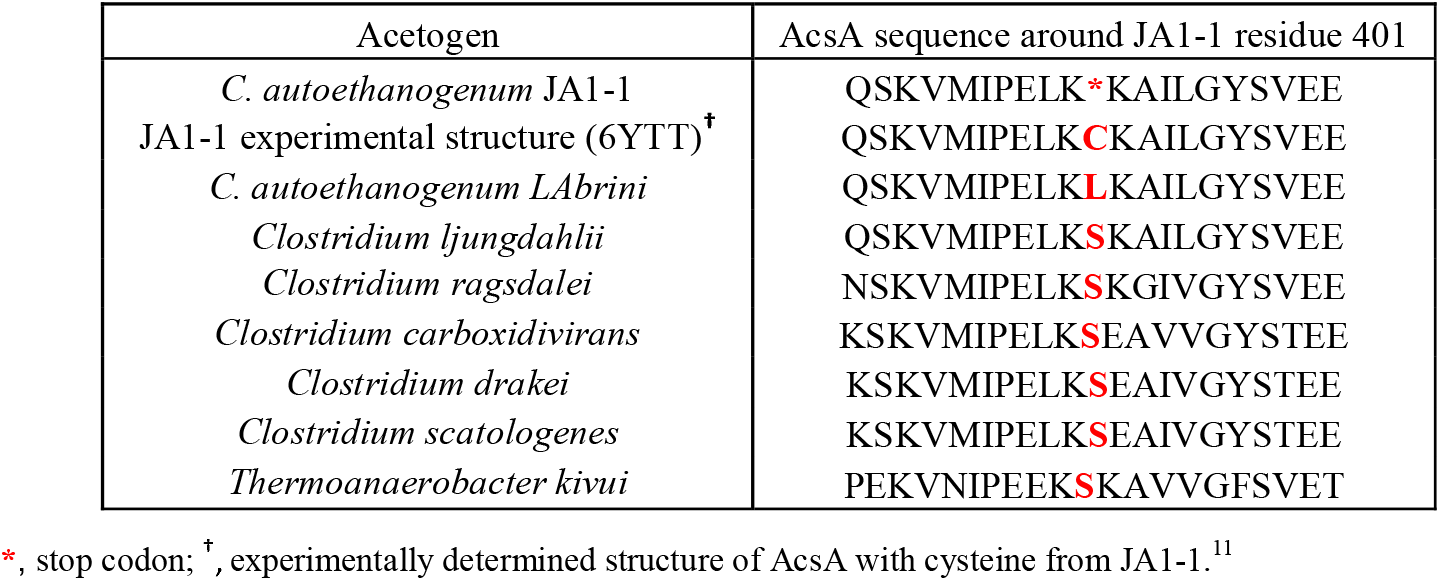
Protein sequence alignment of AcsA (CAETHG_1620-1621) around residue 401 in *Clostridium autoethanogenum* JA1-1 with homologs from closely related acetogens.

AcsA is the only CODH encoded by the C1-fixing gene cluster,^13^ likely since AcsA forms the critical bifunctional enzyme complex with the acetyl-CoA synthase (AcsB or ACS; LABRINI_07965), another cluster member, coupling reduction of CO_2_ to CO with acetyl-CoA formation.^11^ Absolute proteome quantification during *C. autoethanogenum* growth on CO, CO_2_+H_2_, and CO+CO_2_+H_2_ suggests an adapting nature of CODH/ACS activity, shifting between CO oxidation and acetyl-CoA synthesis depending on H_2_ availability, without changes in AcsA expression.^14^ Importantly, the monofunctional CODH CooS1 is significantly more abundant than AcsA on all three gas mixes, and its expression is reduced during ALE of *C. autoethanogenum* on CO_2_+H_2_,^15^ highlighting the relevance of adjusting CooS1 expression for optimal growth. Importantly, inactivation of *cooS1* in *C. autoethanogenum* JA1-1 increases the ethanol-to-acetate ratio on CO and doubles biomass levels on CO_2_+H_2_ with no apparent lag phase, while *cooS2* inactivation had no effects on growth and by-products in batch cultures with a yeast extract-containing medium.^12^ The latter is not surprising as *cooS2* transcript levels are negligible^16–18^ and protein levels remain below the limit of quantification^14,15^ across autotrophic growth of various *C. autoethanogenum* strains. To better understand the role of *cooS1*, it would be relevant to quantify the effects of its deletion also on gas uptake and transcriptome expression, preferably in controlled bioreactor cultures and minimal medium.

In this study, we investigated the importance of the *acsA* internal stop codon by replacing it with leucine or serine, and the effect of *cooS1* deletion in *C. autoethanogenum*. Stop codon removal in *acsA* substantially affected the growth and metabolic by-product profiles during autotrophic batch and continuous cultivation, while deletion of *cooS1* had minimal effects. The latter was consistent with limited transcriptional changes in Δ*cooS1*, while transcriptomics suggested potential transcriptional adjustments linked to reduced robustness and altered product profiles of Leu_SNP. Our findings broaden the understanding of the role and regulation of CODHs in autotrophy and suggest targets for rational engineering of acetogen cell factories.

## EXPERIMENTAL SECTION

### Bacterial strains, growth media, and culture conditions

*Escherichia coli* strains, NEB Turbo and NEB Express (New England Biolabs), were used as hosts for cloning, plasmid construction, and propagation. *E. coli* strains were routinely cultivated in Lysogeny Broth (LB) or LB agar plates supplemented with 100 μg/mL of ampicillin. The *C. autoethanogenum* strain JA1◻ (DSM 10061)^19^ and LAbrini (DSM 115981)^13^ were used as the base strains to create single-nucleotide modifications in *acsA* and deletion of the monofunctional CODH *cooS1*, respectively (see below). Transformants of *C. autoethanogenum* strains were selected on YTF agar containing 10 g/L yeast extract (YE), 16 g/L tryptone, 5 g/L fructose,^20^ and 12 g/L agar, and supplemented with 4 μg/mL clarithromycin whenever required. For autotrophic batch cultivations, strains were grown in a chemically defined PETC-MES medium^21^ without YE with 0.4 g/L of cysteine-HCl·H_2_O as the reducing agent. Bottle headspace was pressurised to 140 kPa with either CO (60% CO and 40% Ar; AS Eesti AGA) or with a syngas mixture (50% CO, 20% CO_2_, 20% H_2_, 10% Ar; AS Eesti AGA).

Batch experiments were conducted in 250 mL Schott bottles containing 50 mL of the liquid medium and incubated horizontally at 37°C with orbital shaking at 120 RPM under strictly anaerobic conditions. Growth was monitored by measuring culture optical density at 600 nm (OD_600_) with frequent sampling during the exponential phase for determining maximum growth rate (μ_max_) and production yields (mmol of product/gram of dry cell weight [gDCW]) for acetate, ethanol, and 2,3-butanediol (2,3-BDO). μ_max_ was calculated using 4-to-6 data points from the exponential phase, resulting in a correlation coefficient of R^2^ ≥ 0.97 (except 0.47, 0.70, 0.74 for acetate for one bio-replicate of Δ*cooS1* on CO, Leu_SNP on syngas, and Ser_SNP on CO, respectively, and 0.67 for ethanol for one bio-replicate of Leu_SNP on syngas) between cultivation time and natural logarithm (ln) of culture OD_600_. Product yields for acetate, ethanol, and 2,3-BDO were calculated during exponential growth by linear regression between the respective product (mmol/L) and biomass concentrations (gDCW/L).

Autotrophic chemostat fermentations (total 19, see Table 2) were carried out as described before.^13,16^ Δ*cooS1* was grown on CO (60% CO and 40% Ar) or syngas (50% CO, 20% CO_2_, 20% H_2_, 10% Ar), Leu_SNP on CO, and Ser_SNP on syngas in a chemically defined medium (without YE) under strictly anaerobic conditions at 37°C and at pH 5 (maintained by 5 M NH_4_OH) with a dilution rate of 1 day^−1^ (D = 1 day^-1^) or 0.5 day^−1^ for Ser_SNP. Chemostat cultivation was performed in 1.4 L Multifors bioreactors (Infors AG) at a working volume of 750 mL and the bioreactors were connected to a Hiden HPR-20-QIC mass spectrometer (Hiden Analytical) for online high-resolution off-gas analysis. Steady-state results were collected after OD, gas uptake, and production rates had been stable (<10% variability) for at least three working volumes. Carbon recoveries and balances were determined as described before.^16^

**Table 2.** Summary of 19 steady-state chemostat cultures with key parameters.

| Strain | Feed gas | Gas flow rate (mL/min) | Agitation (RPM) | D (day<sup>-1</sup>) | Biomass concentration (gDCW/L) | Number of bio-replicates |
| --- | --- | --- | --- | --- | --- | --- |
| LABriini | CO | 50 | 675 | 0.97 ± 0.02 | 1.74 ± 0.06 | 4 |
| Leu_SNP | CO | 50 | 675 | 1.00 ± 0.05 | 1.42 ± 0.03 | 4 |
| Ser_SNP | Syngas | 28 | 575 | 0.50 ± 0.01 | 1.05 ± 0.18 | 4 |
| $\Delta cooS1$ | CO | 50 | 675 | 0.97 ± 0.02 | 1.71 ± 0.08 | 3 |
| $\Delta cooS1$ | Syngas | 50 | 675 | 1.05 ± 0.06 | 1.55 ± 0.10 | 4 |

D, dilution rate; gDCW, gram of dry cell weight. Data are average ± standard deviation between bio-replicates. Syngas composition – 50% CO, 20% CO<sub>2</sub>, 20% H<sub>2</sub>, 10% Ar.

### Genetic engineering

CODH mutant strains with deletion of *cooS1* (CAETHG_3005; LABRINI_15015) in LAbrini and single-nucleotide modifications of *acsA* (CAETHG_1620-1621; LABRINI_08025) in JA1–1 were constructed using CRISPR/nCas9-aided homologous recombination. Details of our genetic engineering workflow, including designing and selecting sgRNAs, plasmid construction and electroporation, colony screening, plasmid curing, and confirmation of transformants are described before.^13,21^ Primers used in this study are in Table S1 while plasmids and strains used or constructed are in Table S2.

### Biomass concentration and extracellular metabolite analysis

Biomass concentration (gDCW/L) was estimated by measuring OD_600_ and using the correlation coefficient of 0.23 between OD_600_ and DCW, as previously established.^22^ Extracellular metabolite analysis was performed using HPLC as described before.^13^

### Bioreactor off-gas analysis

Bioreactor off-gas analysis was performed as described before^16^ to determine the specific gas uptake (CO and H_2_) and production rates (CO_2_ and ethanol) (mmol/gDCW/day). The Faraday Cup detector monitored the intensities of H_2_, CO, ethanol, H_2_S, Ar, and CO_2_ at 2, 14, 31, 34, 40, and 44 amu, respectively.

### Analysis of the AcsA protein structure

UCSF ChimeraX (version 1.9) software^23^ was used for the visualisation, organisation, and manipulation of the experimentally determined atomic structure of AcsA from *C. autoethanogenum* JA1◻1 (PDB ID: 6YU9).^11^ For the substitution of residue 401 with either leucine or serine, the rotamer with the highest probability and the fewest clashes was selected from the ChimeraX rotamer library. Atomic distances between residue 401 and C-cluster residues (C294, C295, C333, C451, C481, C523, and H259) were calculated using CB atom of residue 401 and SG or ND1 for C-cluster residues. Conformational differences between wild-type AcsA (PDB ID: 6YU9)^11^ and mutant structures were analysed by overlay using Matchmaker GUI with default backbone alignment parameters.

### Transcriptome analysis

Transcriptome analysis comparing Leu_SNP (4 bio-replicates on CO) and Δ*cooS1* (4 bio-replicates on CO and 3 on syngas) with LAbrini (4 bio-replicates on CO and 4 on syngas; syngas samples are from chemostats described before^21^) was performed from steady-state chemostat cultures using RNA sequencing (RNA-seq). Sampling, sample preparation, sequencing, and data analysis are described before.^18^ Reads were mapped to the genome of LAbrini (GenBank CP110420.1). Genes were determined to be differentially expressed (DEG) by fold-change > 1.5 and q-value < 0.05 after false discovery rate correction (FDR)^24^ (Table S4).

## RESULTS AND DISCUSSION

### Expression of full-length *acsA* gene product is beneficial for growth in autotrophic batch cultures

AcsA is the indispensable CODH for autotrophy in *C. autoethanogenum*.^12^ We thus aimed to test the effects of two potentially relevant *acsA* SNPs that would replace the stop codon at position 401 with either leucine or serine in the wild-type *C. autoethanogenum* JA1-1 strain (see Introduction). These mutations were targeted in JA1-1 through single-nucleotide base substitutions using a CRISPR/nCas9-aided approach.^13^ Briefly, two sgRNAs were first identified that could potentially realise the target mutations. Next, mutations were incorporated into homology arms, allowing for the replacement of the original stop codon in the genome with target mutations upon successful homologous recombination and DNA cleavage. After plasmid electroporation, transformant colonies were successfully obtained with both sgRNAs; however, only colonies from one sgRNA contained the desired SNPs, albeit mixed with JA1-1 genotype for serine mutants, necessitating an additional round of selective plating to isolate clean SNP strains (Figure S1). Strains were cured and designated as Leu_SNP and Ser_SNP, representing JA1-1 SNP strains with leucine or serine codon replacing the original stop codon at AcsA position 401, respectively. We first characterised growth and by-product profiles of the SNP strains in autotrophic batch cultures using either syngas (50% CO, 20% CO_2_, 20% H_2_) or CO (60% CO) in PETC-MES medium without yeast extract (YE). We compare growth data with the parental strain JA1-1 (grown with YE as it cannot grow without it) and both growth and by-product profiles with strain LAbrini (LAbrini data are from previous works).^13,21^

Both Leu_SNP and Ser_SNP strains showed significantly faster growth compared to JA1-1 on both gases (Figure 1A). While LAbrini and Leu_SNP strains share the same mutation in AcsA at position 401, the latter showed 43% (p-value = 0.001) and 46% (p-value = 0.005) lower maximum specific growth rates (μ_max_) on CO and syngas, respectively (Figure 1A). Leu_SNP also exhibited a longer lag phase and higher variability between bio-replicates compared to LAbrini on both gases (Figure S2). This indicates that the four additional mutations LAbrini acquired during ALE contribute to its superior growth.^13^ For instance, reverse engineering of one of the five mutations, deletion of the sporulation transcriptional activator (*spo0A*), in the starting strain of ALE (JA1-1) recovered LAbrini phenotypes.^13^ Interestingly, Ser_SNP showed growth characteristics similar to LAbrini on both gases, with μ_max_ values of 0.1 ± 0.002 on CO (p-value = 0.3) and 0.074 ± 0.003 on syngas (p-value = 0.9), and thereby grew faster than Leu_SNP (Figure 1A). Notably, both SNP strains and LAbrini grew faster on CO compared to syngas (Figure 1A), suggesting improved CO assimilation resulting from AcsA mutations at position 401.

**Figure 1.**
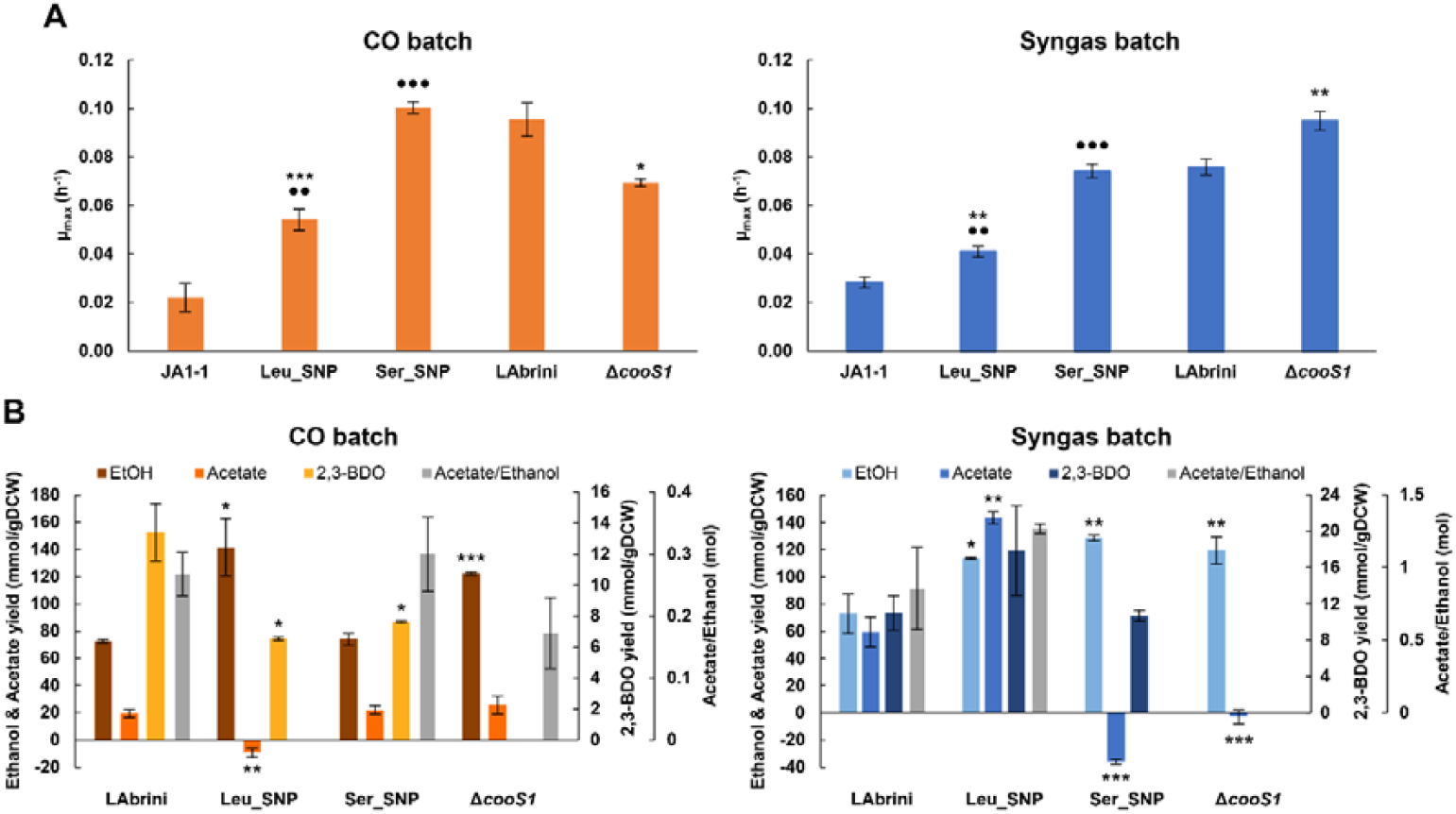
Characterisation of Δ*cooS1* and AcsA SNP strains (Leu_SNP and Ser_SNP) in autotrophic batch conditions. **(A)** Maximum specific growth rates (μ_max_) on CO and syngas without YE, except for JA1, which was grown with YE. Data are average ± standard deviation between three bio-replicates, except for Δ*coos1* on CO and Leu_SNP on syngas, which are with two bio-replicates. **(B)** Growth by-product yields (mmol/gDCW) and acetate/ethanol (Ace/EtOH; mol/mol) on CO and syngas. Acetate/ethanol shown for cultures with simultaneous production of acetate and ethanol (see main text for details). Data are average ± standard deviation between two bio-replicates, except for strains Δ*coos1* and Ser_SNP on syngas that are from three bio-replicates. Dot denotes values statistically different between AcsA SNP strains and JA1-1, while asterisk denotes values statistically different between CODH mutant strains and LAbrini, according to Student’s t-test: • or *, p-value < 0.05; •• or **, p-value < 0.01; ••• or ***, p-value < 0.001. JA1-1 and LAbrini data are from previous work.^13,21^ LAbrini, *C. autoethanogenum* ALE isolate; YE, yeast extract; gDCW, gram of dry cell weight; Ace, acetate; EtOH, ethanol; 2,3-BDO, 2,3-butanediol.

Growth by-product analysis during exponential growth showed remarkably distinct profiles for the two gases (all by-product comparisons use yields; mmol/gDCW). Interestingly, Leu_SNP on CO exhibited a notable 95% increase (p-value = 0.04) in ethanol compared to LAbrini (Figure 1B) with simultaneous consumption of acetate accumulated prior to the exponential phase. The same was seen for Ser_SNP on syngas: 76% higher ethanol production than LAbrini (p-value = 0.003) with simultaneous acetate consumption (Figure 1B). Acetate consumption during autotrophic exponential growth is unusual, as simultaneous production of acetate and ethanol is typically observed. Simultaneous acetate consumption and increased ethanol production indicate elevated levels of reduced ferredoxin (Fd_red_), driving the acetate-to-ethanol conversion via acetaldehyde-ferredoxin oxidoreductase (AOR) activity to maintain redox balance.^25^ On syngas, Leu_SNP also showed higher yields than LAbrini – 141% for acetate (p-value = 0.002) and 56% for ethanol (p-value = 0.03) – likely due to improved assimilation of CO in combination with H_2_, providing extra reducing power (Figure 1B). While acetate and ethanol production for Ser_SNP were comparable to LAbrini on CO, both SNP strains displayed reduced 2,3-BDO yields: 51% for Leu_SNP (p-value = 0.03) and 38% for Ser_SNP (p-value = 0.05) (Figure 1B). Collectively, autotrophic batch cultures showed that Ser_SNP rather than Leu_SNP (which shares AcsA SNP with LAbrini) exhibited growth characteristics and by-product profiles similar to LAbrini on both gases (Figure 1). Additionally, these results show that changing a single amino acid in a key enzyme (AcsA) can significantly impact autotrophic phenotypes in *C. autoethanogenum* JA1-1. This is consistent with reverse engineering a single SNP (acquired by JA1-1 during ALE) in a two-component transcriptional regulator (CLAU_1957), leading to improved JA1-1 phenotypes in our previous study.^26^

### Deletion of *cooS1* leads to metabolic rearrangements in autotrophic batch cultures

Differential expression of the monofunctional CODH CooS1 under various growth conditions (see Introduction) motivated us to investigate its condition-dependent functional importance for autotrophic growth of *C. autoethanogenum*. Markerless and full deletion of *cooS1* was successfully achieved in the *C. autoethanogenum* LAbrini strain using another CRISPR/nCas9-aided approach (Figure S3).^21^ Growth and by-product profiles of Δ*cooS1* were first studied in autotrophic batch cultures using either syngas or CO and PETC-MES medium without YE and compared with the parental strain, LAbrini (LAbrini data are from previous works).^13^,^21^

Δ*cooS1* exhibited a longer lag phase on CO compared to LAbrini (Figure S2) and grew 27% slower (p-value = 0.02) on CO but 28% faster (p-value = 0.007) on syngas than LAbrini (Figure 1A). A previous study also observed slower growth for *cooS1* deletion on CO compared to the parental wild-type JA1-1 strain.^12^ Growth by-products analysis during exponential growth showed that Δ*cooS1* produced significantly more ethanol compared to LAbrini: 69% on CO (p-value = 0.001) and 64% on syngas (p-value = 0.01) (Figure 1B). Strikingly, acetate production was not observed on syngas for Δ*cooS1* (Figure 1B), indicating a higher selectivity for reducing acetate to ethanol in the presence of sufficient reducing power (CO and H_2_). In contrast, acetate production for Δ*cooS1* on CO did not differ from LAbrini (Figure 1B), which contradicts the previously seen lower acetate yield for *cooS1* deletion.^12^ Production of 2,3-BDO was not detected for Δ*cooS1* on either gas.

### Expression of full-length *acsA* gene product affects growth and by-product profiles in autotrophic chemostat cultures

While batch cultures enable the determination of μ_max_ and growth by-product profiles, continuous chemostat cultivation facilitates the generation of high-quality steady-state data^27^ and is the industrially relevant cultivation mode for acetogen gas fermentation.^3^ Therefore, we further characterised all engineered strains and LAbrini in autotrophic chemostats on syngas or CO at a dilution rate (D) of 1 or 0.5 (Ser_SNP) day^-1^ on minimal medium and obtained 19 steady-state datasets (Table 2; LAbrini data on syngas are from a previous work).^13^

The AcsA SNP strains were studied on one gas: Leu_SNP on CO and Ser_SNP on syngas as batch cultures for these combinations showed more distinct by-product profiles compared to LAbrini (Figure 1B). Achieving steady-state conditions was challenging as both Leu_SNP and Ser_SNP cultures exhibited high sensitivity to increasing gas-to-liquid mass transfer during the biomass build-up phase, indicating reduced robustness in terms of bioreactor operation (Figure S4).

Thus, biomass build-up was slow and more time was needed to reach steady-state conditions compared to LAbrini.

Strikingly, Leu_SNP on CO showed significant differences in growth by-product production compared to LAbrini at D = 1 day^-1^: 65% lower specific acetate production rate (q_ace_; mmol/gDCW/day) (p-value < 0.001), 74% higher specific ethanol production rate (q_EtOH_; mmol/gDCW/day) (p-value < 0.001), and 169% higher specific 2,3-BDO production rate (q_2,3-BDO_; mmol/gDCW/day) (p-value < 0.001) (Figure 2A), reflected also in the carbon balances (Figure 2C). Ethanol and acetate production were similarly affected in CO batch cultures (Figure 1B). Furthermore, the ∼5-fold lower Ace/EtOH (0.26 ± 0.03 vs 1.29 ± 0.05 mol/mol for LAbrini) was accompanied by a 27% higher ratio of specific CO_2_ production rate (q_CO2_; mmol/gDCW/day) to specific CO uptake rate (q_CO_; mmol/gDCW/day), q_CO2_/q_CO_ (p-value < 0.001) (Figure 2B). This is consistent with both theoretical stoichiometric calculations^28–30^ and *C. autoethanogenum* steady-state data at various biomass levels^16^ showing that a lower Ace/EtOH leads to higher CO carbon loss as CO_2_. One could speculate that the higher CO uptake by Leu_SNP (p-value = 0.008) (Figure 2B) causes over-supply of Fd_red_ that drives higher production of the reduced by-products ethanol and 2,3-BDO to maintain redox balance since the oxidoreductases within their synthesis pathways from acetyl-CoA, AOR and pyruvate:ferredoxin oxidoreductase (PFOR), respectively, consume Fd_red_ in *C. autoethanogenum*.^16^ Although the by-product profiles for Leu_SNP in both batch and chemostat cultures are attractive for gas fermentation, lower μ_max_ and robustness in bioreactor operation still need to be addressed, potentially with some of the complementary mutations LAbrini acquired during ALE.^13^

**Figure 2.**
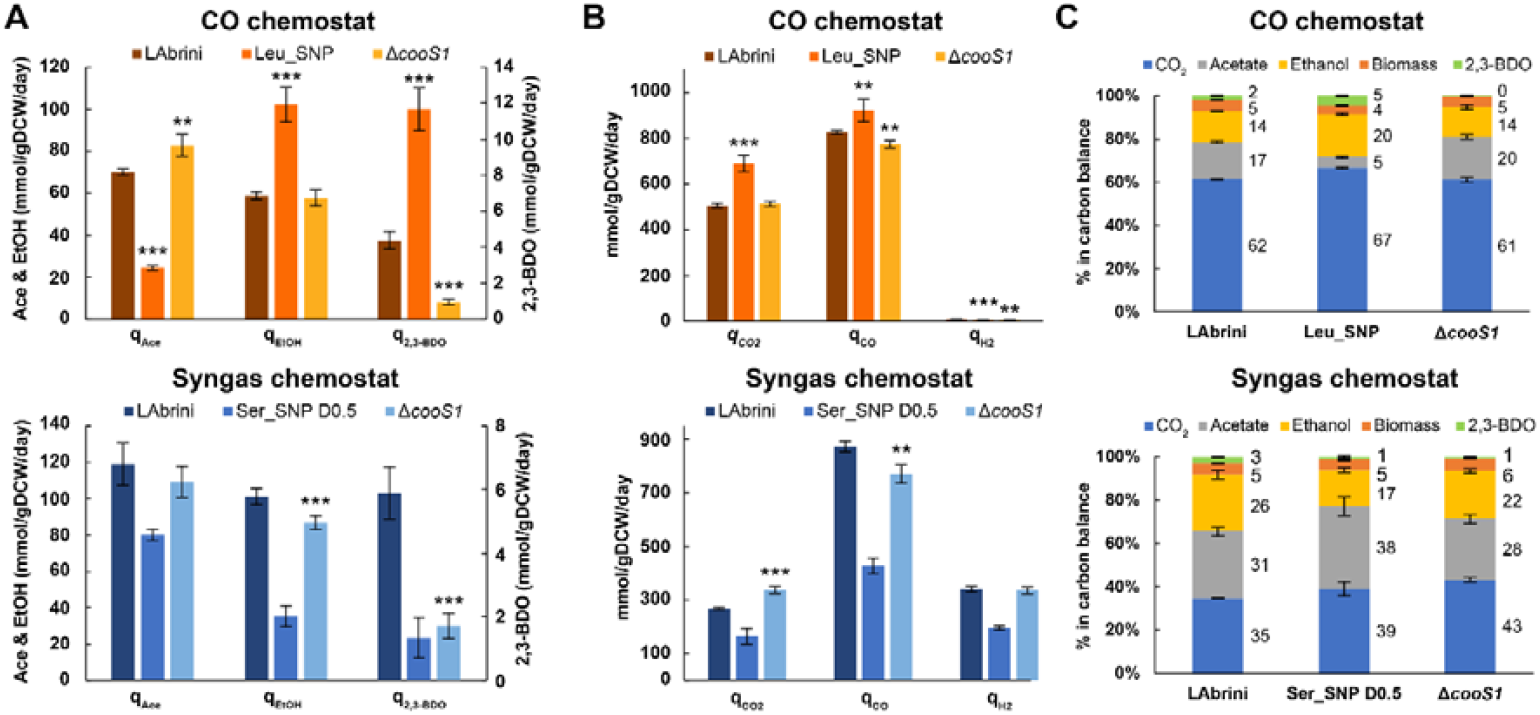
Characterization of Δ*cooS1*, Leu_SNP, Ser_SNP, and LAbrini in steady-state autotrophic chemostat cultures. (**A**) Specific by-product production rates (mmol/gDCW/day) on CO and syngas. (**B)** Specific gas uptake (CO, H_2_ for syngas) and production (CO_2_, H_2_ for CO) rates (mmol/gDCW/day) on CO and syngas. (**C**) Carbon balances. Carbon recoveries were normalised to 100% to have a fair comparison of carbon distributions between different strains. All chemostat data are from dilution rate (D) 1 day^-1^, except from 0.5 day^-1^ for Ser_SNP (Ser_SNP D0.5). Data are average ± standard deviation between four bio-replicates, except three bio-replicates for Δ*coos1* on syngas. Asterisk denotes values statistically different between Leu_SNP or Δ*cooS1* and LAbrini according to Student’s t-test: *, p-value < 0.05; **, p-value < 0.01; ***, p-value < 0.001. Statistical tests not done for Ser_SNP due to the different D. LAbrini data on syngas are from a previous work^13^ LAbrini, *C. autoethanogenum* ALE isolate; Ace, acetate; EtOH, ethanol; 2,3-BDO, 2,3-butanediol; gDCW, gram of dry cell weight; q, specific rate.

Since Ser_SNP on syngas showed particularly high sensitivity to increasing gas-to-liquid mass transfer during the biomass build-up phase (Figure S4B), we targeted and obtained steady-state at D = 0.5 day^-1^. While Ser_SNP showed higher ethanol yields than LAbrini in syngas batch cultures (Figure 1B), a significantly higher Ace/EtOH ratio (2.57 ± 0.46) was determined for Ser_SNP chemostats compared to LAbrini (1.27 ± 0.17 at D = 1 day^-1^) (Figure 2A). A previous study shows high q_ace_ at lower D, with carbon flux moving from acetate towards solvents as D increases.^18^ Interestingly, the q_CO2_/q_CO_ and q_H2_/q_CO_ ratios for Ser_SNP strain were 26% (p-value = 0.009) and 18% (p-value < 0.001) higher than LAbrini (Figure 2B). These higher ratios indicate increased demand for reducing equivalents, potentially for energy homeostasis through CO and H_2_ oxidation. Indeed, Ser_SNP excretes more CO as CO_2_ (Figure 2C).

### Deletion of *cooS1* has minimal effects on carbon fixation and by-product synthesis in autotrophic chemostat cultures

For Δ*cooS1*, we successfully obtained steady-states at D = 1 day^-1^ on both CO and syngas chemostats and compared it with LAbrini (Table 1; LAbrini data on syngas are from a previous work).^13^ In contrast to batch data where Δ*cooS1* showed markedly higher ethanol yields compared to LAbrini (Figure 1), Δ*cooS1* overall displayed lower production of reduced by-products in autotrophic chemostats, except for ethanol which was similar to LAbrini on CO (Figure 2A). This finding highlights the complexity of metabolic responses across different cultivation modes. Interestingly, q_CO_ was lower by 6% (p-value = 0.003) on CO and 12% (p-value = 0.002) on syngas for Δ*cooS1*, indicating reduced CO oxidation activity (Figure 2B). The syngas chemostat cultures of Δ*cooS1* were also characterised by a 26% higher (p-value < 0.001) q_CO2_ compared to LAbrini (Figure 2B), leading to a significant 43% increase (p-value < 0.001) in the q_CO2_/q_CO_ ratio. These data are consistent with the proposition that CooS1 is mainly active in the direction of CO_2_ reduction.^12^ While ethanol production remained similar on CO, an 18% increase (p-value = 0.005) in q_ace_ was observed for Δ*cooS1* on CO, resulting in an increased Ace/EtOH ratio (1.54 ± 0.15 vs 1.29 ± 0.07 for LAbrini, p-value = 0.03) (Figure 2A). Whereas on syngas, Ace/EtOH was not different from LAbrini (1.36 ± 0.15 vs 1.27 ± 0.17 for LAbrini, p-value = 0.5), though 15% lower q_EtOH_ (p-value = 0.002) and 12% higher q_H2_/q_CO_ was observed for Δ*cooS1* (0.44 ± 0.005 vs 0.39 ± 0.01; p-value < 0.001). Expectedly, both Δ*cooS1* and LAbrini diverted more carbon into liquid by-products on syngas compared to CO (Figure 2C).

In a previous study, deletion of *cooS1* in *Thermoanaerobacter kivui* completely eliminated growth on CO.^31^ Although CooS1 is not essential for autotrophic growth of *C. autoethanogenum*, based on this and a previous study,^12^ its deletion impacted growth and metabolism with variable outcomes between batch and chemostat cultures (Figure 1 and 2). Thus, highlighting the relevance of characterising acetogen strains in industrially relevant continuous cultures, which could also inform strain engineering strategies different from batch studies.

### Structural analysis of wild-type AcsA and SNP mutants reveals a conserved structural architecture

The presence of a stop codon in the essential enzyme AcsA raises intriguing questions regarding its effect on enzyme activity and regulation. Both truncated (44 kDa) and full-length (69 kDa) translational products^12^ and peptides across^14^ AcsA are expressed in *C. autoethanogenum*, with the truncated product being expressed at a higher level.^12^ However, the full-length product is very likely the key one, as the C-terminal region after the stop codon contains the ACS binding domain and C-cluster residues critical for AcsA function.^11^ Additionally, the crystal structure of the CODH/ACS complex reveals that ACS forms a complex with the full-length AcsA protein, incorporating a cysteine residue at position 401, which corresponds to the stop codon.^11^ To study the effects of the SNPs we introduced into AcsA, we performed *in silico* mutagenesis at position 401 of the published atomic model of AcsA (PDB: 6YU9) using ChimeraX.^23^ Mutating cysteine in the wild-type structure to leucine or serine revealed no structural differences (Figure 3). Although residue 401 sits in the surface-exposed loop of AcsA and does not directly coordinate metal clusters (C or A clusters), it lies adjacent to gas-transfer tunnels and the ACS binding domain.^11^ We thus calculated solvent-accessible surface area (SASA) values for the three cases and found, as anticipated, an increase for leucine (286.6 Å^2^) and a decrease for serine (223.7 Å^2^) compared to cysteine (241.1 Å^2^), reflecting their respective bulky hydrophobic and small polar characteristics, suggesting subtle side-chain rearrangements. Additionally, we calculated the distance between C-cluster residues and position 401 in wild-type and mutant proteins and found no significant differences (Table S3), indicating that the catalytic site remains intact and unaffected. The widespread effects of the two mutations on fermentation characteristics we detected in both batch and chemostat cultures (Figures 1 and 2) could thus potentially arise from altered translational regulation due to the removal of the stop codon (discussed below). The observed growth impairment in continuous cultures for Ser_SNP was surprising, as serine is often used to probe cysteine functions due to its structural similarity, unlike leucine, which could be more disruptive due to its bulkier nature.^32^

**Figure 3.**
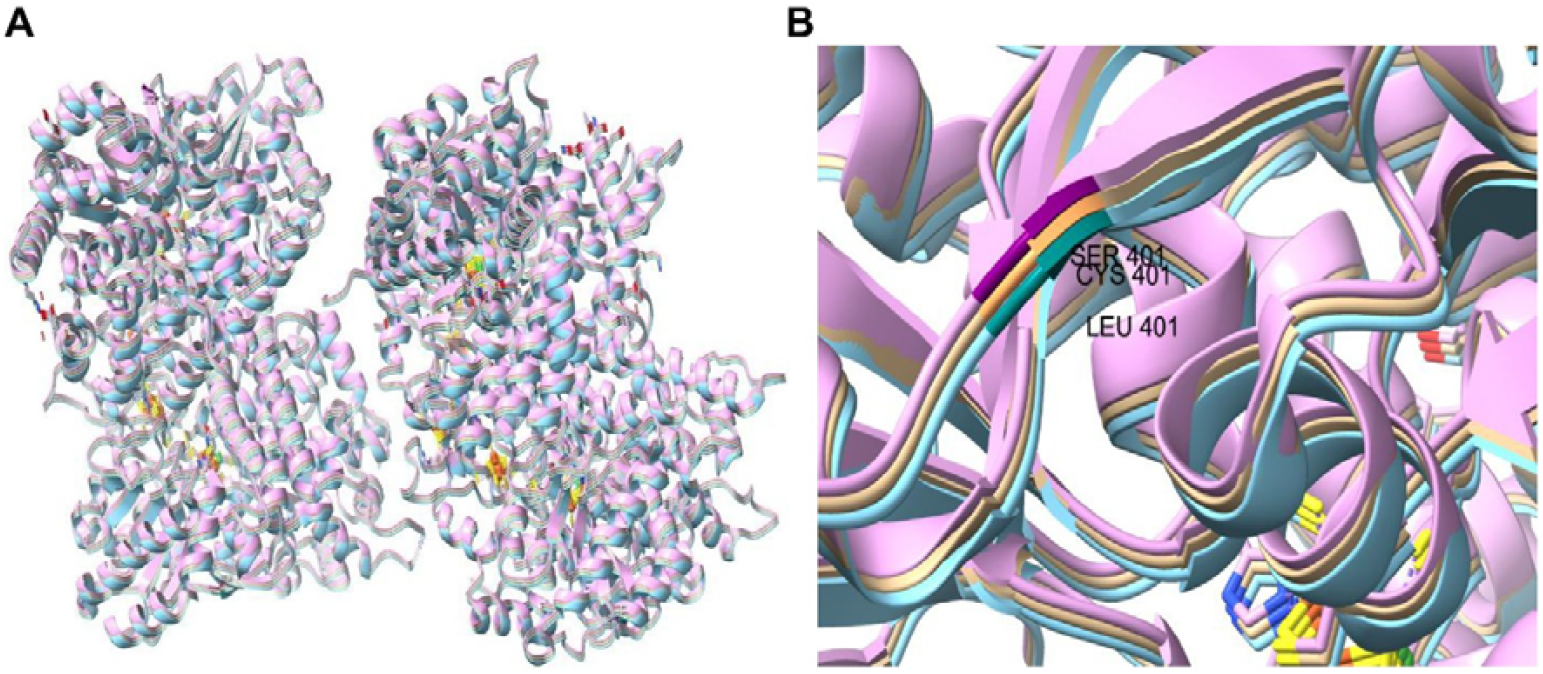
Structural alignment and visualisation of wild-type AcsA and SNP mutant models in ChimeraX. (**A**) Superimposed structure of the AcsA PDB ID:6yu9 (model#1: color tan), Leu_SNP (model#2: color cyan), and Ser_SNP (model#3: color pink). Residue 401 is coloured sandy brown, teal, and purple for the wild-type, Leu_SNP, and Ser_SNP structure, respectively. (**B**) Focused view of the residues surrounding residue 401 in Chain A of the superimposed models.

The benefit of the full-length product is evidenced by the loss of the stop codon in the superior ALE strain LAbrini^13^ and the positive effects of stop codon replacement with leucine on JA1-1 phenotypes in this work. Still, the coexistence of the truncated and full-length products could provide cells with a mechanism to regulate AcsA activity, for instance, to prevent excessive CO_2_ reduction and potential CO toxicity.^33^ Indeed, substantial post-translational modifications of AcsA, particularly in the 25 kDa part following the stop codon (CAETHG_1620), in *C. autoethanogenum* suggest potential regulatory control points.^15^ It is thus not surprising that AcsA has been under evolutionary selection pressure and therefore acquired mutations in several autotrophic ALE experiments with acetogens supporting improved phenotypes.^33^ For instance, an ALE isolate of *Eubacterium limosum* with mutation in AcsA showed improved phenotypes on CO.^34^ Overall, AcsA is an attractive engineering target for improving autotrophic phenotypes.

### Transcriptomics shows minimal gene expression changes in Δ*cooS1* and suggests transcriptional adjustments linked to Leu_SNP phenotype

Lastly, we quantified transcriptomes using RNA sequencing (RNA-seq) in chemostats to investigate, firstly, whether *cooS1* deletion has similarly minimal effects on gene expression as on carbon fixation and by-product synthesis, and secondly, whether differential gene expression could explain the reduced robustness of Leu_SNP compared to LAbrini in terms of bioreactor operation. The latter could be relevant towards improving Leu_SNP robustness for exploiting its attractive product profile for industrial gas fermentation.

High reproducibility between bio-replicate transcriptomes reflected in both bio-replicate clustering and high Pearson correlation coefficients (Figure S5). *cooS1* deletion had limited effects on gene expression on either gas, with 161 differentially expressed genes (DEGs; fold-change > 1.5 and a q-value < 0.05) on syngas, and 182 DEGs on CO relative to LAbrini, of which 66 genes were shared between the two gases (Figure 4A; Table S4). On the contrary, Leu_SNP showed 1,525 DEGs compared to LAbrini (Figure 4B; Table S4), suggesting that the superior robustness of LAbrini for bioreactor operation is realised through extensive transcriptional adjustments modulated by the four additional mutations it carries vs. Leu_SNP.^13^ Notably, Leu_SNP (on CO) and Δ*cooS1*-CO DEGs showed shared enrichment for the KEGG Orthology functional categories pyruvate metabolism (upregulated) and WLP (downregulated) (Table S5).

**Figure 4.**
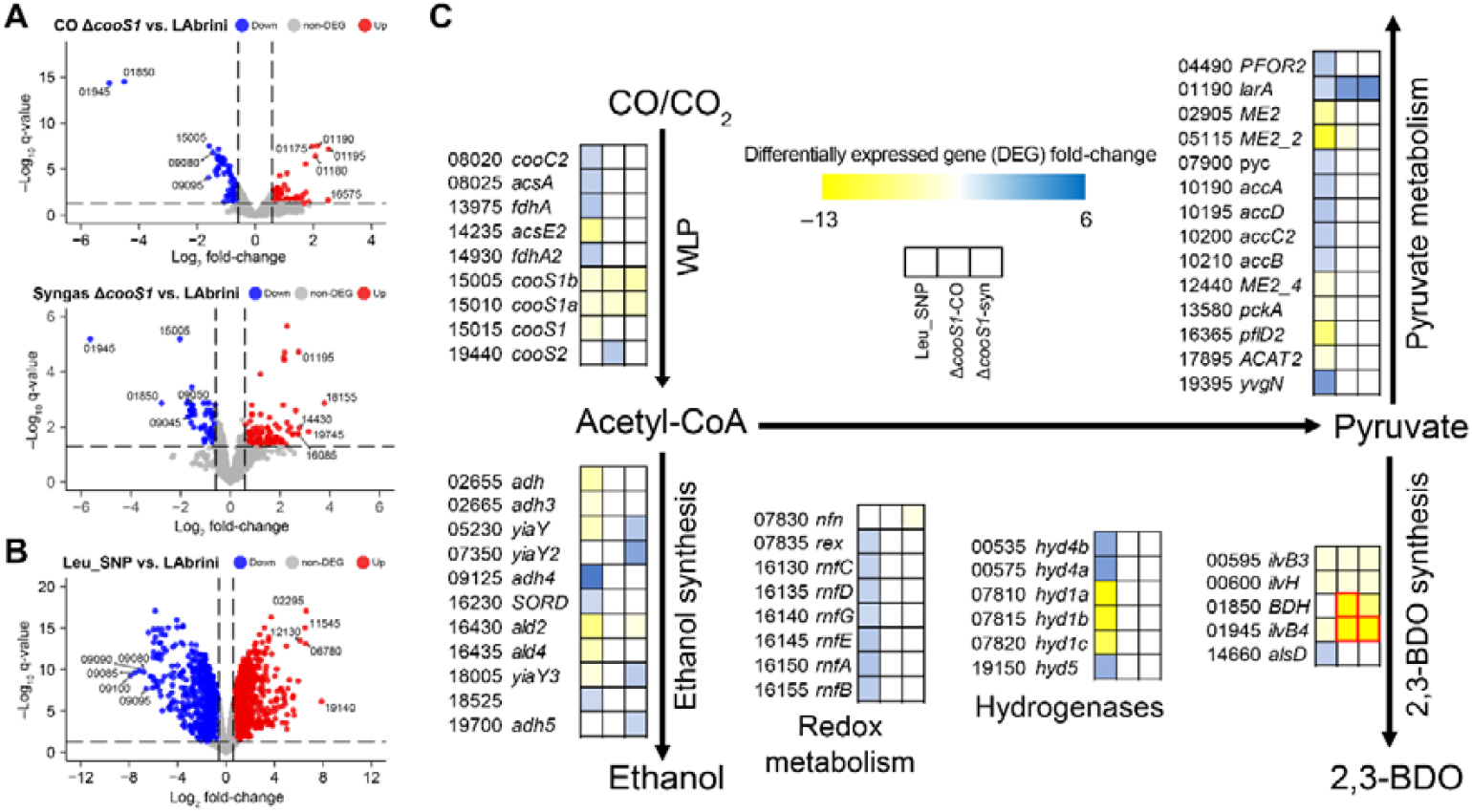
Transcriptional expression differences between Δ*cooS1* and Leu_SNP with LAbrini steady-state autotrophic chemostat cultures. (**A**) Volcano plots showing differentially expressed genes (DEGs) between Δ*cooS1* and LAbrini. **(B**) Volcano plot showing DEGs between Leu_SNP and LAbrini. Top five up- and down-regulated DEGs are indicated by LAbrini gene IDs preceded with ‘LAbrini_’. Dashed lines denote DEG threshold (fold-change > 1.5 and q-value < 0.05). (**C**) DEGs in acetogen central metabolism. Genes that are DEGs (fold-change > 1.5 and q-value < 0.05) at least in one comparison are shown. Data with fold-changes < ◻3 are highlighted with red borders. Non-DEG data have white boxes. For each gene, LAbrini gene IDs, preceded with ‘LAbrini_’, and gene names^14^ are shown. See legend on figure for sample comparison info (all strains vs. LAbrini) and heatmap scale. See Table S4 for DEG data and gene descriptions. WLP, Wood-Ljungdahl pathway; 2,3-BDO, 2,3-butanediol.

Expectedly, we could not detect *cooS1* (LABRINI_15015) transcript expression in Δ*cooS1* (Figure 4C). Interestingly, Δ*cooS1* displayed 2-to-4-fold downregulation of CooS1 subunits (15005, 15010) independent of the gas substrate (Figure 4A and C). CooS1 and its subunits were also downregulated (∼2-fold) in Leu_SNP. While *acsA* (08025) expression was unchanged in Δ*cooS1*, it was upregulated 2-fold in Leu_SNP along with the neighbouring CODH accessory protein CooC2 (08020), presumably involved in CODH’s active site maturation,^35,36^ potentially contributing to increased q_CO_ in Leu_SNP (Figure 2B). The differential expression of CooS1 components with unchanged *acsA* that we detected in Δ*cooS1* aligns with previous observations in *C. autoethanogenum*.^14,37^ Importantly, *cooS2* (19440) was ∼2-fold upregulated in Δ*cooS1* cultures grown on CO, but not on syngas, suggesting a compensatory transcriptional response for CO metabolism in the absence of *cooS1*.

The reduced robustness of Leu_SNP in terms of bioreactor operation could potentially arise from compromised CO fixation through the WLP. Indeed, we detected a 5-fold repression of an alternative methyl-tetrahydrofolate (THF) cobalamin methyltransferase (14235) in Leu_SNP that could be supporting the main methyl-THF cobalamin methyltransferase AcsE (07970) in the transfer of the methyl group from methyl-THF to the corrinoid iron-sulfur proteins AcsC (07975) and AcsD (07980), which in turn supply ACS with the methyl group for acetyl-CoA synthesis in the WLP.^5^ These potential adverse effects on the WLP from 14235 repression could, however, have been complemented by 2-fold upregulation of both formate dehydrogenase isozymes (13975 and 14930) in Leu_SNP. Notably, transcriptional regulation of WLP components appears to be coordinated with hydrogenases, as multiple were differentially expressed in Leu_SNP (Figure 4C; Table S4).

Links between transcriptional regulation of specific by-product synthesis pathway genes and respective by-product profiles remain elusive in acetogens.^1^ Studies in *C. autoethanogenum* suggest this can be due to the dominant post-translational regulation of metabolic fluxes.^14,25,38^ While multiple genes associated with ethanol production were downregulated in Leu_SNP (Figure 4C), the most abundant alcohol dehydrogenase – Adh4 (09125) – in *C. autoethanogenum*^14^ was 6-fold upregulated, potentially realising the higher q_EtOH_ (Figure 2A). At the same time, AORs were not differentially regulated in Leu_SNP, which is consistent with the expected post-translational regulation of its acetate to acetaldehyde conversion.^17^ Similarly, the greatly lower q_Ace_ in Leu_SNP (Figure 2A) was not realised through repression of acetate synthesis genes. Regarding 2,3-BDO synthesis, while repression of 2,3-BDO pathway genes correlated with lower q_2,3-BDO_ for Δ*cooS1*, this was not the case for Leu_SNP (Figure 2A and 4C). In Δ*cooS1*-syn data, iron-containing alcohol dehydrogenases (05230, 07350, and 18005) were upregulated 2-to-4-fold (Figure 4C) while q_EtOH_ was lower (Figure 2A).

Notably, Leu_SNP shifts in growth and product profiles coincided with significant transcriptional rewiring in redox metabolism. Firstly, the entire multi-subunit Fd-NAD^+^ oxidoreductase Rnf complex (16130◻6155), which generates the proton motive force in *C. autoethanogenum* to drive the ATPase,^39,40^ was upregulated 2-to-3-fold in Leu_SNP (Figure 4C). This is consistent with Leu_SNP being the only strain with higher q_CO_ (i.e. extra supply of Fd_red_ for Rnf) and with higher q_EtOH_ (i.e. extra supply of NADH from Rnf) (Figure 2). Higher supply of Fd_red_ from elevated q_CO_ in Leu_SNP could also explain the upregulation of PFOR 04490, consuming Fd_red_ and leading to increased q_2,3-BDO_ (Figure 4C). Secondly, 97-fold upregulation of an NAD(P)-dependent oxidoreductase (02295) could contribute to shifts in redox metabolism in Leu_SNP (Table S4). Lastly, potentially relevant is also the strong up-to 109-fold repression of the gene cluster 09000□09080) associated with ethanolamine utilisation bacterial-microcompartments (BMC) (Figure 4C; Table S4). Transcriptional regulation of the BMC cluster has been detected before,^18^ while its relevance to *C. autoethanogenum* autotrophy remains unclear.

## CONCLUSIONS

Our findings demonstrate a close functional association between CODHs and acetogen metabolism, with genetic modifications to CODHs markedly influencing *C. autoethanogenum* physiology. Distinct phenotypic outcomes observed in *acsA* SNP strains, guided by adaptive evolutionary and comparative genomic evidence, indicate that single-nucleotide missense substitutions in *acsA* can modulate function and regulation without altering enzyme structure. In contrast, the limited phenotypic changes in the *cooS1* deletion strain reveal its condition-dependent role in autotrophy and highlight functional asymmetry among CODHs. While transcriptomics showed limited transcriptional changes in Δ*cooS1*, it suggested potential transcriptional adjustments linked to reduced robustness and altered product profiles of Leu_SNP, which need to be explored further in future work. Overall, this work broadens our understanding of CODH function and regulation in acetogens and illustrates how evolutionary insights can inform rational engineering of acetogens.

## AUTHOR CONTRIBUTIONS

Conceptualization: KMS, KR, and KV; Methodology: KMS, KR, PRP, CV, and KV; Formal analysis: KMS, KR, PRP, and KV; Investigation: KMS, KR, PRP, and CV; Resources: KV; Writing – Original Draft: KMS and KV; Writing – Review & Editing: KMS, KR, PRP, CV, and KV; Supervision: KV; Project Administration: KV; Funding Acquisition: KV.

## SUPPORTING INFORMATION

Results of Sanger sequencing for SNP strains, growth profiles of CODH mutant strains and LAbrini, PCR plus gel electrophoresis for Δ*coos1*, bioreactor operational differences between SNP strains and LAbrini, transcriptome data reproducibility (DOCX); additional experimental details with lists of primers, strains, and plasmids; results for inter-residue distance calculation using ChimeraX, transcriptome analysis for differentially expressed genes (DEGs), enrichment analysis for DEGs (XLSX).

## ACKNOWLEDGEMENTS

We thank fellow group members Ugochi Jennifer Nwaokorie, Nilesh Sheshrao Kolhe, and Victoria Chinonyerem Udemezue for their feedback on the manuscript. This work was funded by the European Union’s Horizon 2020 grant agreement N810755 and co-funded by the European Union and Ministry of Education and Research via project TEM-TA104. The research was conducted using the research infrastructure “National Centre for Translational and Clinical Research”, funded by the Estonian Research Council (TARISTU24-TK22).

## REFERENCES

(1) Pavan, M.; Reinmets, K.; Garg, S.; Mueller, A. P.; Marcellin, E.; Köpke, M.; Valgepea, K. Advances in systems metabolic engineering of autotrophic carbon oxide-fixing biocatalysts towards a circular economy. Metab. Eng. 2022, 71 (9), 117–141. 10.1016/j.ymben.2022.01.015.

(2) Nguyen, A. D.; Lee, E. Y. Engineered Methanotrophy: A Sustainable Solution for Methane-Based Industrial Biomanufacturing. Trends Biotechnol. 2021, 39 (4), 381–396. 10.1016/j.tibtech.2020.07.007.

(3) Köpke, M.; Simpson, S. D. Pollution to products: recycling of ‘above ground’ carbon by gas fermentation. Curr. Opin. Biotechnol. 2020, 65 (5), 180–189. 10.1016/j.copbio.2020.02.017.

(4) Liew, F. E.; Nogle, R.; Abdalla, T.; Rasor, B. J.; Canter, C.; Jensen, R. O.; Wang, L.; Strutz, J.; Chirania, P.; De Tissera, S.; Mueller, A. P.; Ruan, Z.; Gao, A.; Tran, L.; Engle, N. L.; Bromley, J. C.; Daniell, J.; Conrado, R.; Tschaplinski, T. J.; Giannone, R. J.; Hettich, R. L.; Karim, A. S.; Simpson, S. D.; Brown, S. D.; Leang, C.; Jewett, M. C.; Köpke, M. Carbon-negative production of acetone and isopropanol by gas fermentation at industrial pilot scale. Nat. Biotechnol. 2022, 40 (3), 335–344. 10.1038/s41587-021-01195-w.

(5) Ragsdale, S. W. Enzymology of the Wood–Ljungdahl Pathway of Acetogenesis. Ann. N. Y. Acad. Sci. 2008, 1125 (1), 129–136. 10.1196/annals.1419.015.

(6) Valgepea, K.; De Souza Pinto Lemgruber, R.; Abdalla, T.; Binos, S.; Takemori, N.; Takemori, A.; Tanaka, Y.; Tappel, R.; Köpke, M.; Simpson, S. D.; Nielsen, L. K.; Marcellin, E. H_2_ drives metabolic rearrangements in gas-fermenting Clostridium autoethanogenum. Biotechnol. Biofuels 2018, 11 (1), 55. 10.1186/s13068-018-1052-9.

(7) Bertsch, J.; Müller, V. CO Metabolism in the Acetogen Acetobacterium woodii. Appl. Environ. Microbiol. 2015, 81 (17), 5949–5956. 10.1128/AEM.01772-15.

(8) Baum, C.; Zeldes, B.; Poehlein, A.; Daniel, R.; Müller, V.; Basen, M. The energy-converting hydrogenase Ech2 is important for the growth of the thermophilic acetogen Thermoanaerobacter kivui on ferredoxin-dependent substrates. Microbiol. Spectr. 2024, 12 (4). 10.1128/spectrum.03380-23.

(9) Bertsch, J.; Müller, V. Bioenergetic constraints for conversion of syngas to biofuels in acetogenic bacteria. Biotechnol. Biofuels 2015, 8 (1), 210. 10.1186/s13068-015-0393-x.

(10) Brown, S. D.; Nagaraju, S.; Utturkar, S.; De Tissera, S.; Segovia, S.; Mitchell, W.; Land, M. L.; Dassanayake, A.; Köpke, M. Comparison of single-molecule sequencing and hybrid approaches for finishing the genome of Clostridium autoethanogenum and analysis of crispr systems in industrial relevant clostridia. Biotechnol. Biofuels 2014, 7 (1), 40. 10.1186/1754-6834-7-40.

(11) Lemaire, O. N.; Wagner, T. Gas channel rerouting in a primordial enzyme: Structural insights of the carbon-monoxide dehydrogenase/acetyl-CoA synthase complex from the acetogen Clostridium autoethanogenum. Biochim. Biophys. Acta - Bioenerg. 2021, 1862 (1), 148330. 10.1016/j.bbabio.2020.148330.

(12) Liew, F.; Henstra, A. M.; Winzer, K.; Köpke, M.; Simpson, S. D.; Minton, N. P. Insights into CO_2_ Fixation Pathway of Clostridium autoethanogenum by Targeted Mutagenesis. MBio 2016, 7 (3), 427–443. 10.1128/mBio.00427-16.

(13) Ingelman, H.; Heffernan, J. K.; Harris, A.; Brown, S. D.; Shaikh, K. M.; Saqib, A. Y.; Pinheiro, M. J.; de Lima, L. A.; Martinez, K. R.; Gonzalez-Garcia, R. A.; Hawkins, G.; Daleiden, J.; Tran, L.; Zeleznik, H.; Jensen, R. O.; Reynoso, V.; Schindel, H.; Jänes, J.; Simpson, S. D.; Köpke, M.; Marcellin, E.; Valgepea, K. Autotrophic adaptive laboratory evolution of the acetogen Clostridium autoethanogenum delivers the gas-fermenting strain LAbrini with superior growth, products, and robustness. N. Biotechnol. 2024, 83, 1–15. 10.1016/j.nbt.2024.06.002.

(14) Valgepea, K.; Talbo, G.; Takemori, N.; Takemori, A.; Ludwig, C.; Mahamkali, V.; Mueller, A. P.; Tappel, R.; Köpke, M.; Simpson, S. D.; Nielsen, L. K.; Marcellin, E. Absolute Proteome Quantification in the Gas-Fermenting Acetogen Clostridium autoethanogenum . mSystems 2022, 7 (2). 10.1128/msystems.00026-22.

(15) Heffernan, J.; Garcia Gonzalez, R. A.; Mahamkali, V.; McCubbin, T.; Daygon, D.; Liu, L.; Palfreyman, R.; Harris, A.; Koepke, M.; Valgepea, K.; Nielsen, L. K.; Marcellin, E. Adaptive laboratory evolution of Clostridium autoethanogenum to metabolize CO_2_ and H_2_ enhances growth rates in chemostat and unravels proteome and metabolome alterations. Microb. Biotechnol. 2024, 17 (4). 10.1111/1751-7915.14452.

(16) Valgepea, K.; de Souza Pinto Lemgruber, R.; Meaghan, K.; Palfreyman, R. W.; Abdalla, T.; Heijstra, B. D.; Behrendorff, J. B.; Tappel, R.; Köpke, M.; Simpson, S. D.; Nielsen, L. K.; Marcellin, E. Maintenance of ATP Homeostasis Triggers Metabolic Shifts in Gas-Fermenting Acetogens. Cell Syst. 2017, 4 (5), 505–515. 10.1016/j.cels.2017.04.008.

(17) Marcellin, E.; Behrendorff, J. B.; Nagaraju, S.; Detissera, S.; Segovia, S.; Palfreyman, R. W.; Daniell, J.; Licona-Cassani, C.; Quek, L. E.; Speight, R.; Hodson, M. P.; Simpson, S. D.; Mitchell, W. P.; Köpke, M.; Nielsen, L. K. Low carbon fuels and commodity chemicals from waste gases – systematic approach to understand energy metabolism in a model acetogen. Green Chem. 2016, 18 (10), 3020–3028. 10.1039/c5gc02708j.

(18) de Lima, L. A.; Ingelman, H.; Brahmbhatt, K.; Reinmets, K.; Barry, C.; Harris, A.; Marcellin, E.; Köpke, M.; Valgepea, K. Faster Growth Enhances Low Carbon Fuel and Chemical Production Through Gas Fermentation. Front. Bioeng. Biotechnol. 2022, 10, 879578. 10.3389/fbioe.2022.879578.

(19) Abrini, J.; Naveau, H.; Nyns, E. J. Clostridium autoethanogenum, sp. nov., an anaerobic bacterium that produces ethanol from carbon monoxide. Arch. Microbiol. 1994, 161 (4), 345–351. 10.1007/BF00303591.

(20) Liew, F.; Henstra, A. M.; Kpke, M.; Winzer, K.; Simpson, S. D.; Minton, N. P. Metabolic Engineering of Clostridium autoethanogenum for Selective Alcohol Production. Metab. Eng. 2017, 40, 104–114. 10.1016/j.ymben.2017.01.007.

(21) Nwaokorie, U. J.; Reinmets, K.; de Lima, L. A.; Pawar, P. R.; Shaikh, K. M.; Harris, A.; Köpke, M.; Valgepea, K. Deletion of genes linked to the C_1_-fixing gene cluster affects growth, by-products, and proteome of Clostridium autoethanogenum. Front. Bioeng. Biotechnol. 2023, 11, 1167892. 10.3389/fbioe.2023.1167892.

(22) Peebo, K.; Valgepea, K.; Nahku, R.; Riis, G.; Õun, M.; Adamberg, K.; Vilu, R. Coordinated activation of PTA-ACS and TCA cycles strongly reduces overflow metabolism of acetate in Escherichia coli. Appl. Microbiol. Biotechnol. 2014, 98 (11), 5131–5143. 10.1007/s00253-014-5613-y.

(23) UCSF ChimeraX Home Page https://www.rbvi.ucsf.edu/chimerax.

(24) Benjamini, Y.; Hochberg, Y. Controlling the False Discovery Rate: A Practical and Powerful Approach to Multiple Testing. J. R. Stat. Soc. Ser. B 1995, 57 (1), 289–300. 10.1111/J.2517-6161.1995.TB02031.X.

(25) Mahamkali, V.; Valgepea, K.; de Souza Pinto Lemgruber, R.; Plan, M.; Tappel, R.; Köpke, M.; Simpson, S. D.; Nielsen, L. K.; Marcellin, E. Redox controls metabolic robustness in the gas-fermenting acetogen Clostridium autoethanogenum. Proc. Natl. Acad. Sci. U. S. A. 2020, 117 (23), 13168–13175. 10.1073/pnas.1919531117.

(26) Ingelman, H.; Shaikh, K. M.; Valgepea, K. Reverse Engineered Gas Fermenting Acetogen Strains Recover Enhanced Phenotypes From Autotrophic Adaptive Laboratory Evolution. Microb. Biotechnol. 2025, 18 (8), e70208. 10.1111/1751-7915.70208.

(27) Adamberg, K.; Valgepea, K.; Vilu, R. Advanced continuous cultivation methods for systems microbiology. Microbiol. (United Kingdom) 2015, 161 (9), 1707–1719. 10.1099/mic.0.000146.

(28) Bertsch, J.; Müller, V. Bioenergetic constraints for conversion of syngas to biofuels in acetogenic bacteria. Biotechnol. Biofuels 2015, 8 (1), 210. 10.1186/s13068-015-0393-x.

(29) Liew, F. M.; Martin, M. E.; Tappel, R. C.; Heijstra, B. D.; Mihalcea, C.; Köpke, M. Gas Fermentation-A Flexible Platform for Commercial Scale Production of Low-Carbon-Fuels and Chemicals from Waste and Renewable Feedstocks. Front. Microbiol. 2016, 7, 694. 10.3389/fmicb.2016.00694.

(30) Molitor, B.; Richter, H.; Martin, M. E.; Jensen, R. O.; Juminaga, A.; Mihalcea, C.; Angenent, L. T. Carbon recovery by fermentation of co-rich off gases – Turning steel mills into biorefineries. Bioresour. Technol. 2016, 215 (8), 386–396. 10.1016/j.biortech.2016.03.094.

(31) Jain, S.; Katsyv, A.; Basen, M.; Müller, V. The monofunctional CO dehydrogenase CooS is essential for growth of Thermoanaerobacter kivui on carbon monoxide. Extremophiles 2021, 26 (1), 4. 10.1007/s00792-021-01251-y.

(32) Qiu, H.; Honey, D. M.; Kingsbury, J. S.; Park, A.; Boudanova, E.; Wei, R. R.; Pan, C. Q.; Edmunds, T. Impact of cysteine variants on the structure, activity, and stability of recombinant human α-galactosidase A. Protein Sci. 2015, 24 (9), 1401–1411. 10.1002/pro.2719.

(33) Davin, M. E.; Thompson, R. A.; Giannone, R. J.; Mendelson, L. W.; Carper, D. L.; Martin, M. Z.; Martin, M. E.; Engle, N. L.; Tschaplinski, T. J.; Brown, S. D.; Hettich, R. L. Clostridium autoethanogenum alters cofactor synthesis, redox metabolism, and lysine-acetylation in response to elevated H_2_:CO feedstock ratios for enhancing carbon capture efficiency. Biotechnol. Biofuels Bioprod. 2024 171 2024, 17 (1), 119-. 10.1186/s13068-024-02554-w.

(34) Kang, S.; Song, Y.; Jin, S.; Shin, J.; Bae, J.; Kim, D. R.; Lee, J. K.; Kim, S. C.; Cho, S.; Cho, B. K. Adaptive Laboratory Evolution of Eubacterium limosum ATCC 8486 on Carbon Monoxide. Front. Microbiol. 2020, 11, 402. 10.3389/fmicb.2020.00402.

(35) Carlson, E. D.; Papoutsakis, E. T. Heterologous Expression of the Clostridium carboxidivorans CO Dehydrogenase Alone or Together with the Acetyl Coenzyme A Synthase Enables Both Reduction of CO_2_ and Oxidation of CO by Clostridium acetobutylicum. Appl. Environ. Microbiol. 2017, 83 (16), 829–846. 10.1128/AEM.00829-17.

(36) Gregg, C. M.; Goetzl, S.; Jeoung, J. H.; Dobbek, H. AcsF Catalyzes the ATP-Dependent Insertion of Nickel into the Ni,Ni-[4Fe4S] Cluster of Acetyl-CoA Synthase. J. Biol. Chem. 2016, 291 (35), 18129–18138. 10.1074/jbc.M116.731638.

(37) Heffernan, J.; Garcia Gonzalez, R. A.; Mahamkali, V.; McCubbin, T.; Daygon, D.; Liu, L.; Palfreyman, R.; Harris, A.; Koepke, M.; Valgepea, K.; Nielsen, L. K.; Marcellin, E. Adaptive laboratory evolution of Clostridium autoethanogenum to metabolize CO_2_ and H_2_ enhances growth rates in chemostat and unravels proteome and metabolome alterations. Microb. Biotechnol. 2024, 17 (4), e14452. 10.1111/1751-7915.14452.

(38) Greene, J.; Daniell, J.; Köpke, M.; Broadbelt, L.; Tyo, K. E. J. Kinetic ensemble model of gas fermenting Clostridium autoethanogenum for improved ethanol production. Biochem. Eng. J. 2019, 148, 46–56. 10.1016/j.bej.2019.04.021.

(39) Hess, V.; Gallegos, R.; Jones, J. A.; Barquera, B.; Malamy, M. H.; Müller, V. Occurrence of ferredoxin:NAD+ oxidoreductase activity and its ion specificity in several Gram-positive and Gram-negative bacteria. PeerJ 2016, 4, e1515. 10.7717/peerj.1515.

(40) Tremblay, P.; Zhang, T.; Dar, S. A.; Leang, C.; Lovley, D. R. The Rnf Complex of Clostridium Ljungdahlii Is a Proton-Translocating Ferredoxin:NAD+ Oxidoreductase Essential for Autotrophic Growth. MBio 2012, 4 (1), e00406–12. 10.1128/mBio.00406-12.

