## Supporting figures for "Genetic engineering of carbon monoxide dehydrogenases produces distinct autotrophic phenotypes in *Clostridium autoethanogenum*"

Kurshedaktar Majibullah Shaikh^1,#^, Kristina Reinmets^1,#^, Pratik Rajendra Pawar^1^,

Clara Vida G. C. Carneiro^1^, Kaspar Valgepea^1,*^

^1^Institute of Bioengineering, University of Tartu, Nooruse 1, Tartu, 50411, Estonia

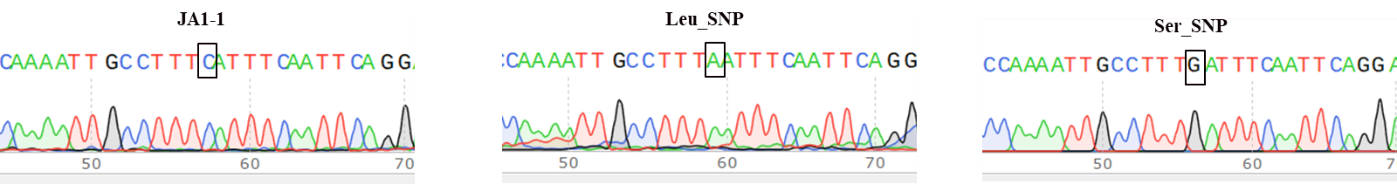

**Figure S1.** **Sanger sequencing chromatogram confirming the targeted single-nucleotide mutation at position 401 of *acsA* (box).**

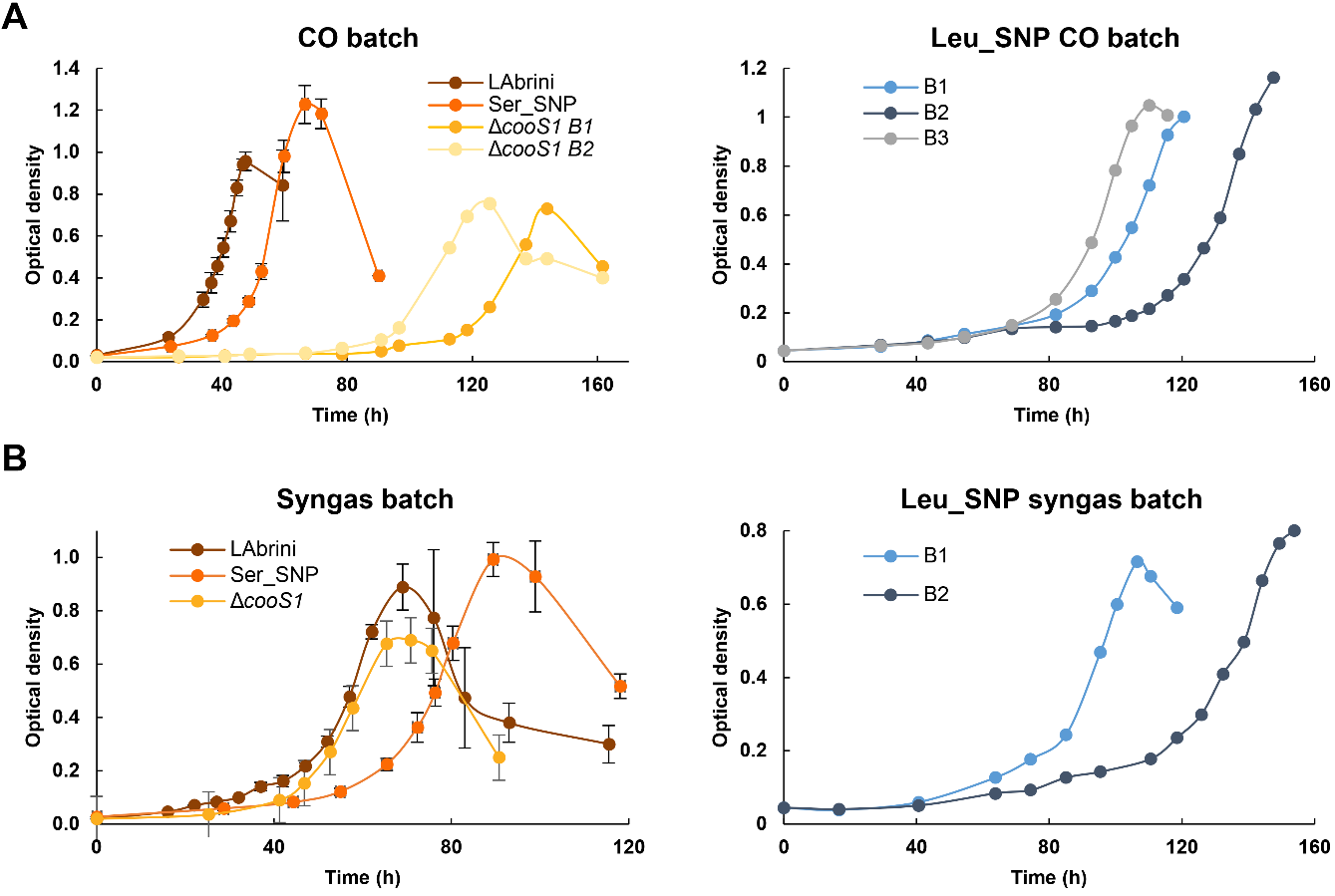

**Figure S2.** **Autotrophic bottle batch growth of Δ*cooS1,* Leu_SNP, and Ser_SNP strains in PETC-MES without YE.** (**A**) Growth on CO; (**B**) Growth on syngas. Data represent average ± standard deviation between three bio-replicates, except for Δ*coos1* on CO and Leu_SNP for which independent bio-replicate data are shown. B, bio-replicate.

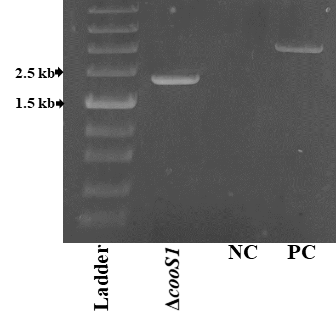

**Figure S3.** **PCR confirmation of Δ*cooS1* strain.** PC, positive control (Non-transformed LAbrini); NC, negative control (no template)

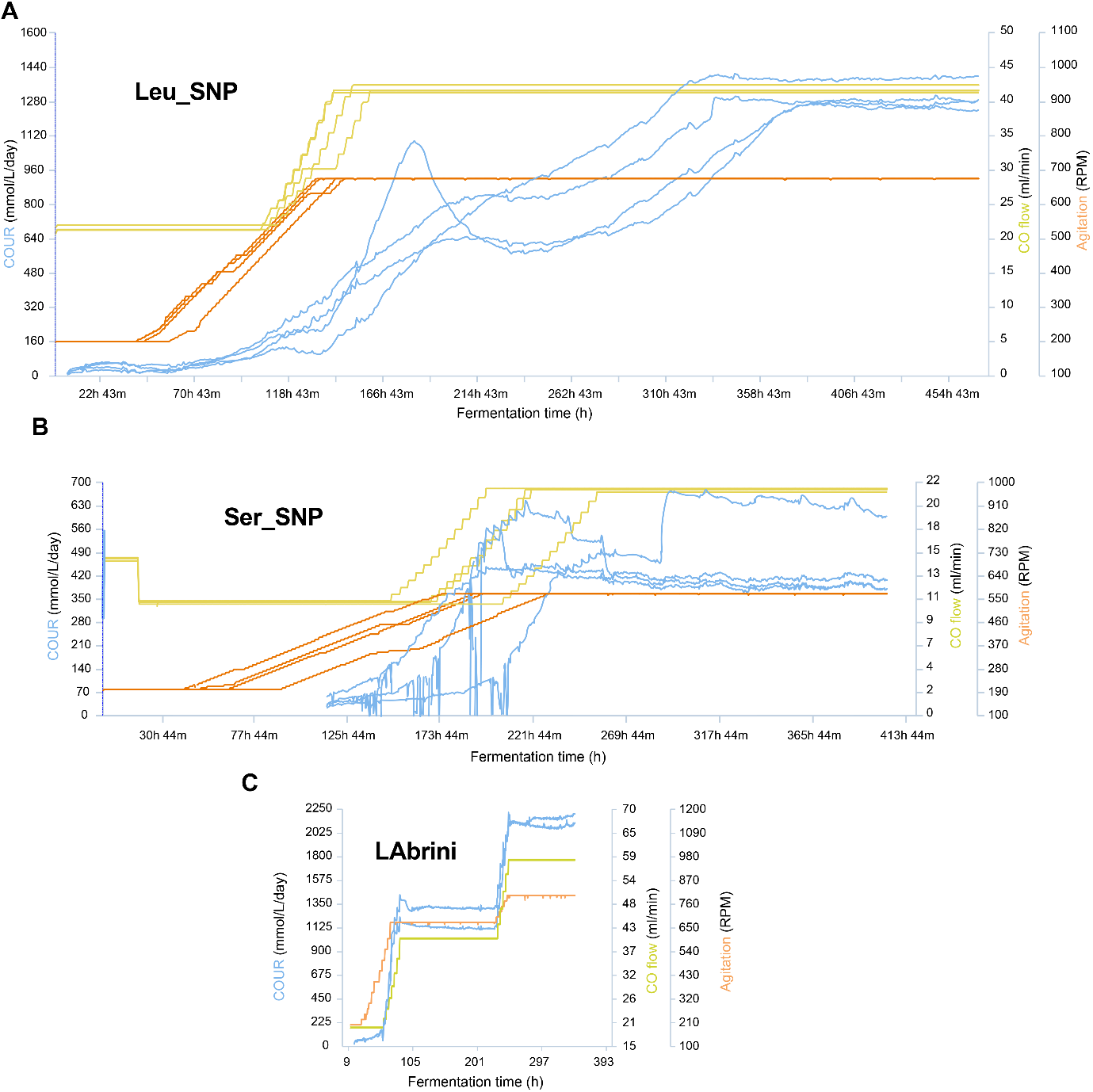

**Figure S4. Reduced robustness of SNP strains in terms of bioreactor operation for chemostat cultivations.** (**A**) Leu_SNP on CO at dilution rate (D) 1 day^-1^ with four bio-replicates. (**B**) Ser_SNP on syngas at D = 0.5 day^-1^ with four bio-replicates. COUR data before ~110h missing due to MS malfunction. (**C**) LAbrini data on syngas are for two of the four bio-replicates that achieved steady-states at D = 1 and 2 day^-1^ (Figure 4 in Ingelman et al. 2024; doi: 10.1016/J.NBT.2024.06.002; reproduced with permission from authors). COUR, CO uptake rate (mmol/L/day); RPM, revolutions per minute.

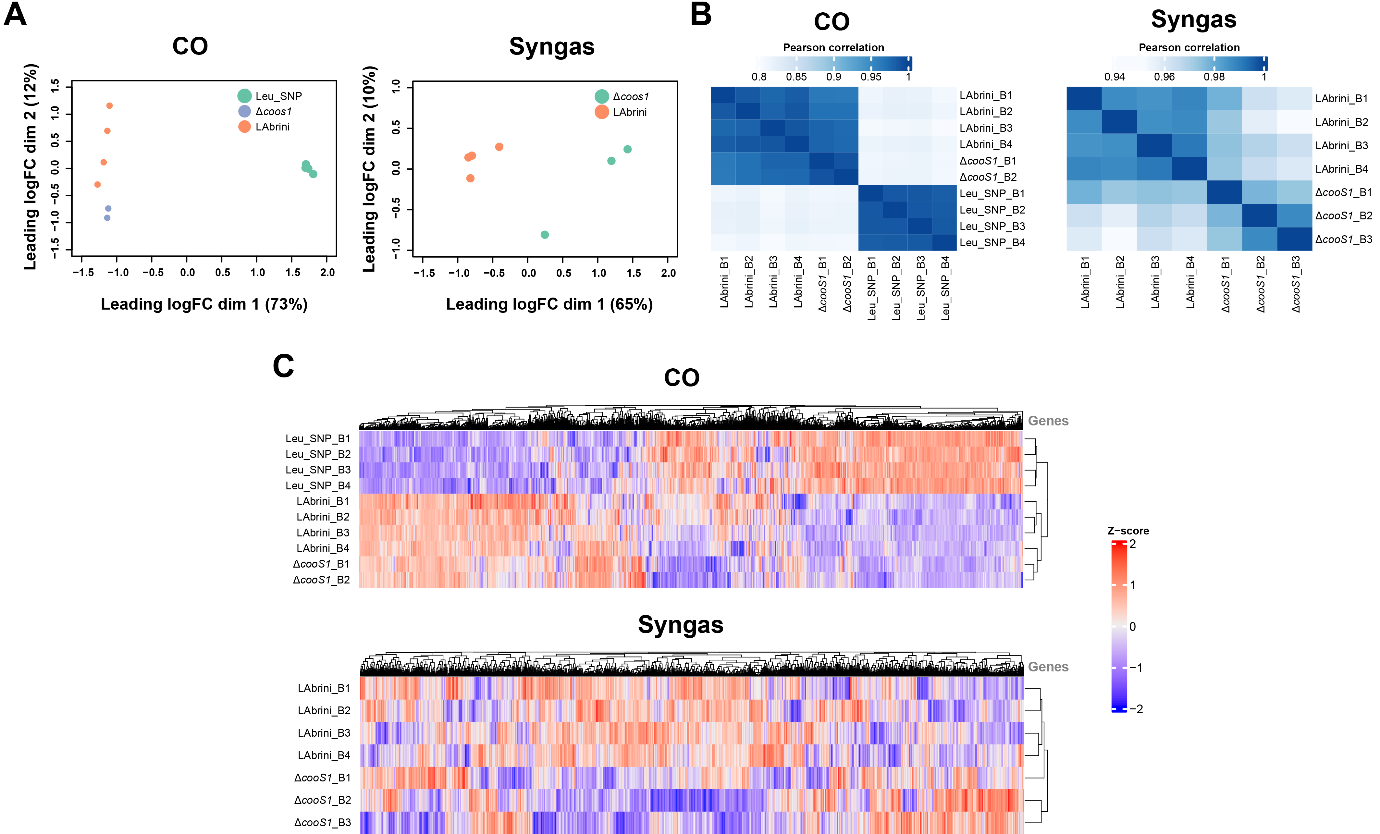

**Figure S5. Transcriptome data reproducibility in chemostats.** (**A**) Multidimensional scaling (MDS) of bio-replicate transcript expression changes. The leading log-fold-change (logFC) between a pair of samples is defined as the root-mean-square average of the top largest log_2_-fold-changes between those two samples. (**B**) Pearson correlations between bio-replicate transcript abundances (RPKM). (**C**) Hierarchical clustering of bio-replicate transcript abundances (Z-scores based on RPKMs). RPKM, reads per kilobase of transcript per million mapped reads. B, bio-replicate.
